# Diversity of Ganglion Cell Responses to Saccade-like Image Shifts in the Primate Retina

**DOI:** 10.1101/2022.08.12.503725

**Authors:** Steffen Nitsche, Mohammad H. Khani, Dimokratis Karamanlis, Yunus C. Erol, Sören J. Zapp, Matthias Mietsch, Dario A. Protti, Fernando Rozenblit, Tim Gollisch

**Affiliations:** Department of Ophthalmology, University Medical Center Göttingen, Göttingen, Germany; Bernstein Center for Computational Neuroscience Göttingen, Göttingen, Germany; International Max Planck Research School for Neurosciences, Göttingen, Germany; Laboratory Animal Science Unit, German Primate Center, Göttingen, Germany; German Center for Cardiovascular Research, Partner Site Göttingen, Göttingen, Germany; School of Medical Sciences (Neuroscience), The University of Sydney, Sydney, NSW, Australia; Cluster of Excellence “Multiscale Bioimaging: from Molecular Machines to Networks of Excitable Cells” (MBExC), University of Göttingen, Göttingen, Germany

## Abstract

Saccades are a fundamental part of natural vision. They interrupt fixations of the visual gaze and rapidly shift the image that falls onto the retina. These stimulus dynamics can cause activation or suppression of different retinal ganglion cells, but how they affect the encoding of visual information in different types of ganglion cells is largely unknown. Here, we recorded spiking responses to saccade-like shifts of luminance gratings from ganglion cells in isolated marmoset retinas and investigated how the activity depended on the combination of pre- and post-saccadic images. All identified cell types, On and Off parasol and midget cells as well as a type of Large Off cells, displayed distinct response patterns, including particular sensitivity to either the pre- or the post-saccadic image or combinations thereof. In addition, Off parasol and Large Off cells, but not On cells, showed pronounced sensitivity to whether the image changed across the transition. Stimulus sensitivity of On cells could be explained based on their responses to step changes in light intensity, whereas Off cells, in particular, parasol and the Large Off cells, seem to be affected by additional interactions that are not triggered during simple light-intensity flashes. Together, our data show that ganglion cells in the primate retina are sensitive to different combinations of pre- and post-saccadic visual stimuli. This contributes to the functional diversity of the retina’s output signals and to asymmetries between On and Off pathways and provides evidence of signal processing beyond what is triggered by isolated steps in light intensity.

## Introduction

Research in visual neuroscience is often motivated by features of natural stimuli, like contrast, color, and dynamics of the visual environment. Fully natural stimuli, however, are so rich in features that elicited neural responses can often be difficult to interpret. It has thus proven useful to focus on a particular feature of natural stimuli and recreate it artificially in abstract form, like changes in brightness investigated with uniform grayscale stimuli (Hartline, 1938), spatial structure at varying resolution with gratings (Enroth-Cugell and Robson, 1966), or small objects of interest with moving spots of light (Lettvin et al., 1959). A specific feature that dominates much of the dynamics of vision is given by saccades—rapid eye or body movements that shift the point of fixation and occur multiple times per second in humans (Yarbus, 1967). Saccades strongly structure the visual stimuli that fall onto the retina and thereby shape the neural signals sent from the eye to the rest of the brain.

Retinal ganglion cells can display diminished responsiveness during saccades or saccade-like image shifts, which is thought to contribute to saccadic suppression—the phenomenon of reduced visual perception around the time of saccades (Roska and Werblin, 2003; Wurtz, 2008; Idrees et al., 2020). However, psychophysical studies have found that saccade kinematics can be re-adjusted during the saccade if the target is moved at saccade onset (Gaveau et al., 2003), indicating that meaningful visual processing must occur during saccades. Indeed, suppression does not affect all retinal ganglion cells and instead cells can exhibit various responses to and during saccades (Noda and Adey, 1974; Amthor et al., 2005; Sivyer et al., 2019), and responses after saccade offset may furthermore be particularly informative about the newly fixated image (Segev et al., 2007; Krishnamoorthy et al., 2017). A study on macaque ganglion cells found responses to natural scenes to be determined by the eye-movement-like temporal structure of the stimulus (Schottdorf and Lee, 2021).

Little is known, however, about how the rapid succession of fixations and brief transitions affect the encoding of visual information, in particular for the primate retina. One hypothesis might be that a saccade acts like a reset, allowing a new, independent snapshot of the visual world after saccade offset. Retinal ganglion cells would then respond to the newly fixated image according to how strongly the newly encountered visual contrast activates their receptive fields. Yet, the offset of the previously fixated image also presents a potent stimulus just a few tens of milliseconds earlier. In principle, ganglion cell activity after a saccade may thus depend in a complex fashion on the combination of pre- and post-saccadic stimulus patterns and be additionally influenced by the image motion during the saccade.

Here, we investigate how primate retinal ganglion cell responses to saccade-like image shifts are shaped by the combination of pre- and post-saccadic stimulation of the receptive field. Based on multielectrode array recordings of ganglion cells from the marmoset retina and functional identification of the major ganglion cell types, we find cell-type-specific differences in the sensitivity to saccadic stimulus features. A model with nonlinear spatial stimulus integration and response properties derived from simple light-intensity flashes could partially capture these response characteristics. We conclude that saccades trigger cell-type-specific signal-processing mechanisms that contribute to functional asymmetries between On and Off ganglion cells and broaden the scope of visual features encoded by the retina across saccades.

## Materials and Methods

### Tissue Preparation and Electrophysiology

We used retinas of three adult marmoset monkeys (*Callithrix jacchus*) of either sex (2 male, 1 female), aged 4, 7, and 15 years. Retinal tissues were obtained immediately after euthanasia from animals used by other researchers, in accordance with national and institutional guidelines and as approved by the institutional animal care committee of the German Primate Center and by the responsible regional government office (Niedersächsisches Landesamt für Verbraucherschutz und Lebensmittelsicherheit, permit number 33.19-42502-04-17/2496).

After enucleation, the eyes were dissected, and the cornea, lens, and vitreous humor were carefully removed to gain direct access to the retina. The tissue was then transferred into a light-tight chamber containing oxygenated (95% O_2_ and 5% CO_2_) Ames’ medium (Sigma-Aldrich, Munich, Germany), supplemented with 6 mM D-glucose, and buffered with 22 mM NaHCO_3_ to maintain a pH of 7.4. After 1-2 hours of dark adaptation, the retina was dissected into smaller pieces. For each recording, a piece of peripheral retina was isolated from the pigment epithelium and transferred to a multielectrode array (MultiChannel Systems, Reutlingen, Germany; either 60 or 252-electrode planar arrays, 30 μm electrode diameter and 100 μm minimum electrode spacing). The preparation was performed under infrared illumination with a stereomicroscope equipped with night-vision goggles. During the recording, the retina was perfused with the oxygenated Ames’ medium (4-5 ml/min), and the temperature of the recording chamber was kept constant around 33°C using an inline heater (PH01, MultiChannel Systems, Reutlingen, Germany) and a heating element below the array. The remaining retina tissue continued to be stored in the light-tight chamber and constantly perfused with oxygenated Ames’ medium for later recordings.

The multielectrode array signals were amplified, band-pass filtered (300 Hz to 5 kHz), and stored digitally at 25 kHz (60-electrode arrays) or 10 kHz (252-electrode arrays), using the software MC-Rack 4.6.2 (MultiChannel Systems). Spike sorting was performed with a modified version of the sorting software Kilosort (Pachitariu et al., 2016), available at https://github.com/MouseLand/Kilosort (original) and https://github.com/dimokaramanlis/KiloSortMEA (modified version). The output of Kilosort was visually inspected and manually curated with the software Phy2 (https://github.com/cortex-lab/phy). Only units with a well-separated cluster of voltage traces and a clear refractory period were included for further analyses.

### Visual Stimulation

Visual stimuli were generated by custom-made software written in C++ and OpenGL and displayed on a gamma-corrected monochromatic white OLED monitor (eMagin) with a refresh rate of 60 Hz and 800 × 600 pixels. The stimuli were projected onto the retina using a telecentric lens (Edmund Optics) resulting in a pixel size of 7.5 μm × 7.5 μm on the retina. All stimuli used in this study had a mean light level of either 1.9 or 3.3 mW/m^2^ in the mesopic to low photopic regime. The same light level was also used for homogeneous illumination between stimuli. Before the start of an experiment, the projection of the stimulus screen was focused on the photoreceptor layer by visual inspection via a microscope.

### Estimation of Receptive Fields, Nonlinearities, and Autocorrelations

In order to characterize the receptive fields of recorded cells and their autocorrelation functions, a spatiotemporal binary white-noise stimulus on a checkerboard layout was presented. Stimulus pixels had a size of 60 μm by 60 μm on the retina (37.5 μm by 37.5 μm for one experiment). Each pixel was updated independently and pseudo-randomly at the monitor refresh rate of 60 Hz to display either black or white (100% Michelson contrast). The stimulus consisted of an alternating sequence of 1500 frames (25 s) of independent, non-repeating white noise and 300 frames (5 s) of a fixed, repeated white-noise sequence. For the present study, only the independent white-noise segments were used. The stimulus was presented for 30-40 minutes leading to about 100,000 frames of independent white noise.

Receptive fields were determined by first calculating the spatiotemporal spike-triggered average (STA) of a cell’s responses to the independent white noise (Chichilnisky, 2001). We used a temporal window of 30 frames (0.5 s) for the STA. To separate the STA into a temporal and a spatial filter, we selected the element (pixel and time point) with the maximum absolute value in the STA after smoothing with a spatial Gaussian filter of 60 μm standard deviation. The temporal filter was then defined as the time course of the selected pixel in the unsmoothed STA and the spatial filter as the corresponding unsmoothed frame. A two-dimensional Gaussian was fitted to the spatial filter. The standard deviations of the Gaussian were reduced to 80% of the original to account for the observation that white-noise stimuli activate the surround less strongly than flashed stimuli and therefore often overestimate the receptive field size relative to more flash-like stimuli with larger spatial structure (Wienbar and Schwartz, 2018). The reduced Gaussian function was normalized to a volume of unity and taken as an estimate of the receptive field, and the effective receptive-field diameter was defined as the diameter of a circle with the same area as the 1.5-sigma ellipse of this Gaussian function. Receptive-field outlines were displayed as this 1.5-sigma ellipse.

To characterize a cell’s contrast–response relationship, we computed the cell’s nonlinearity as part of the linear-nonlinear model (Chichilnisky, 2001). This was done by computing the dot product of every frame in the white-noise stimulus with the cell’s spatial filter and convolving the resulting sequence with its temporal filter to obtain a generator signal for each stimulus frame. To reduce noise, only pixels within in the smallest rectangular window still containing the 3.75-sigma ellipse of the Gaussian were included in this computation. The generator signals were then binned into ten bins with equal number of data points, and the average spike count and generator signal were calculated for each bin. The resulting histogram was fitted with the function *a*·*C*(*b·g+c*), where *C* is the cumulative distribution function of the normal distribution, *g* the generator signal, and *a, b, c* free parameters, and the fit was treated as the cell’s nonlinearity.

Spike-train autocorrelation functions were computed over 50 ms at a resolution of 0.04 ms (25 kHz recordings) or 0.1 ms (10 kHz recording) from the responses to the white-noise stimulus, then smoothed with a Gaussian filter with standard deviation of ten data points, and normalized to a sum of unity.

### Classification of Retinal Ganglion Cells

To be able to investigate whether different cell types play distinct roles during saccades, we first classified cells manually in a way similar to the procedure in Field et al. (2007). For each recording, we computed the first two principal components of all temporal filters. We then constructed scatter plots of the projections of the temporal filters onto the first principal component against the projection onto the second principal component as well as against the effective receptive-field diameter. The scatter plots yielded clustered groups of cells, corresponding to On and Off midget and parasol cells, respectively, and a fifth cell type that we here call Large Off cells. Most recorded cells could readily be assigned to one of the clusters based on these scatter plots. For cells which lay at the borders of clusters, assignment to a cluster was additionally based on examining the spike-train autocorrelation function, the detailed shapes of the temporal filter and the nonlinearity, and the positioning of the receptive field relative to receptive fields of other cells in the nearby clusters. Only few cells could then not clearly be assigned to one of the analyzed types and were thus excluded from further analyses.

### Saccadic Stimulus

To stimulate the retina with saccade-like image shifts, we used a stimulus based on rapid movements of a spatial square-wave grating. The grating had a Michelson contrast of 60% and a bar width of 90 μm on the retina. The stimulus mimicked an alternating sequence of fixations lasting 533 ms each and saccade-like transitions of 67 ms. During each fixation, the grating remained static at one of four equally spaced spatial phases, which are called Positions 1, 2, 3 and 4. The sequence of positions was chosen pseudo-randomly. Transitions moved the grating from one position to the next by translating the grating by about two full grating periods, as previously used in (Krishnamoorthy et al., 2017). Note, however, that in our experiments, these motion transitions were depicted for only four monitor frames such that due to aliasing the screen did not show a smooth movement of the grating but rather a quick succession of various grating positions. For half of the transitions, chosen pseudo-randomly, the transition was masked by a uniform gray screen at mean intensity of the grating. The saccadic stimulus was presented for 12-20 minutes resulting in 1200-2000 transitions.

We analyzed the responses of each cell to the transitions according to the combination of grating position before the saccade, termed starting position, and the grating position after the saccade, termed target position. For each of the resulting 16 combinations of starting and target position, we collected the cell’s responses to calculate a 350-ms-long peri-stimulus time histogram (PSTH) with a bin size of 10 ms. For quantitative analyses of response amplitudes just after the onset of the transition and after the onset of the new fixation, we divided each PSTH into a first response phase, ranging from 30 ms after onset of the transition until 30 ms (20 ms for one experiment with shorter response latencies) after onset of the target position, and a second response phase ranging from 30 ms (20 ms) to 200 ms after onset of the target position. Peak responses in each phase were then determined after separately smoothing each corresponding PSTH segment with a Gaussian of 20 ms standard deviation, using zero-padding.

We collected the detected peak firing rates in each of the two response phases in two 4×4 response matrices, one for each response phase. These response matrices were then Fourier-transformed (two-dimensional discrete Fourier transform), yielding two complex-valued matrices whose entries quantify the amplitudes and phases of different basic patterns in the response matrices. We took the absolute values of the matrix entries, thereby disregarding the phase and only keeping the amplitude of the patterns. From each transformed matrix, we extracted the three entries (0, 1), (1, 0), and (3, 1), which correspond to specific sensitivities (to target position, to starting position, and to change across the transition) as described in the main text. These three entries were combined in a three-dimensional vector, yielding two vectors for each cell that needed to be normalized in order to make them comparable across cells. This was done by comparing the Frobenius norms of the two Fourier-transformed response matrices and dividing both vectors by the larger one. Intuitively, this relates each specific response modulation pattern to the total response modulation in the response phase with the stronger modulations. After having observed that three out of the six entries of the two vectors are almost always close to zero, the remaining three values were combined into a final sensitivity vector: the (0, 1) component, i.e. starting sensitivity, of the vector of the first response phase, and the (1, 0) and (3, 1) components, i.e. target and change sensitivity, of the vector of the second response phase.

### Responses to Brightness Steps

We used the responses of ganglion cells to full-field uniform steps in light intensity as a basis for modeling their responses to the saccadic stimulus. The light intensity steps lasted 0.5 s, going alternatingly to white (+100% Michelson contrast) and black (−100% Michelson contrast), separated by 1.5 s of gray mean-intensity illumination. This stimulus was repeated for 30-90 cycles taking a total of 2-6 minutes. PSTHs were computed with a bin size of 10 ms both for the entire cycle duration for display in the figures, and for a time window of 400 ms following each of the four changes in light intensity for the modeling of responses to the saccadic stimulus (see below).

### Modeling

We compared the ability of two computational models to predict ganglion cell responses to the saccadic stimulus based on a cell’s responses to the brightness steps. For these analyses, the transition period was modeled as homogeneous gray illumination at mean light intensity, based on the observation that measured responses did not differ between such masked transitions and transitions with shifting grating position.

Both models use a weighted summation of the firing-rate profiles measured under brightness steps. The two models only differed in whether contrast signals over the receptive field were integrated linearly or nonlinearly when computing the weights. For the linear model, we first computed the net change in visual contrast over the receptive field for each transition in the saccade stimulus and used this as a weight for the corresponding response trace as measured under the brightness steps to arrive at the model prediction. Conversely, for the nonlinear model, each pixel essentially contributed to the firing-rate prediction according to its own contrast change, and the averaging over the receptive field only occurred afterwards, thus preventing the cancelation of activity by simultaneous brightening and darkening in the different parts of the receptive field.

Concretely, the two model predictions *R*_linear_(*t*) and *R*_nonlinear_(*t*) of a cell were computed from the cell’s responses *R*_x→y_(*t*) under brightness steps, where x→y stands for the different transitions white→gray (w→g), black→gray (b→g), gray→white (g→w), and gray→black (g→b). For a given transition from the pre-saccadic stimulus 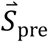 to the post-saccadic stimulus 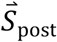, we used:

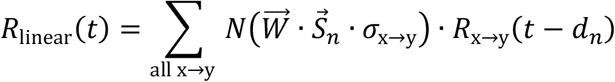

and

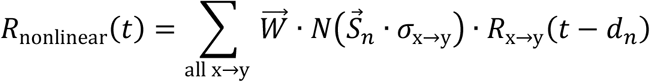

Here, *N*(·) is a thresholding function, setting negative inputs to zero, and 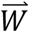 is the (Gaussian) receptive field of the cell, evaluated at the pixel centers and correspondingly denoted as a vector. 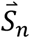 stands for the appropriate pixel-wise stimulus, 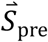 for the pre-saccadic image, used for w→g and b→g, and 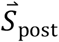 for the post-saccadic image used for g→w and g→b. The elements of the 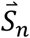 are −1 and 1 for black and white pixels, respectively. The time delay *d*_n_ is used to shift the responses corresponding to the occurrence of 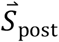 by the transition duration, hence *d*_post_ = 67 ms and *d*_pre_ = 0. The scalar factor σ_x→y_ is used to adjust the sign of the stimulus depending on whether a brightening or darkening is considered and whether the onset or offset of the stimulus pattern is considered; hence σ_x→y_ = 1 for w→g and for g→w and −1 otherwise. For the *R*_x→y_ (*t*), we used the 400 ms long PSTHs of the cell after the full-field brightness step from x to y as calculated in the section “Responses to Brightness Steps” and zero-padded them for time points outside of the 400 ms window.

Finally, for comparison with measured responses to the saccade stimulus, we allowed for a constant latency shift that is applied in the same way to the firing-rate predictions for all 16 transitions. This aims at accounting for the overall faster responses under brightness steps, owing to the higher contrast of this stimulus as compared to the saccade stimulus. We fitted the latency shift for each cell by selecting the shift in the range of 0 to 50 ms with the maximal Pearson correlation between the data and the prediction. For most cells, the latency shift was less than 30 ms. Predictions were calculated for a 350 ms window starting at the onset of the transition, which is the same duration as the PSTHs calculated for the saccadic stimulus. For figures showing predicted responses, we jointly scaled the predictions for the 16 transitions to the same peak value as in the corresponding data from that cell, again to account for differences in applied overall contrast.

We used two measures to evaluate the model performance. Firstly, we compared the response matrices calculated from the predicted responses with the cell’s experimental response matrices using a modified coefficient of determination. The modeled response matrices were scaled to the same mean as the experimental ones to accommodate for the different contrasts. The modified coefficient of determination between one pair of experimental and modeled response matrices was then calculated as

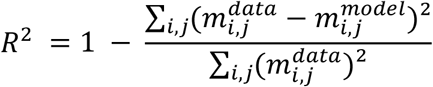

where m_i,j_ are the entries of the response matrix of the experimental data or of the prediction, respectively. This modified coefficient of determination corresponds to substituting the mean of the data’s response matrix in the denominator of the regular coefficient of determination with zero. This was necessary because for cells that responded indifferently to all transitions the mean is already the best prediction for the response matrix. Thus, the null hypothesis rather should be an unresponsive cell with a response matrix consisting of zeros.

Furthermore, we used the Euclidean distance in the three-dimensional sensitivity vector space as a measure for model performance. The distance was computed between the sensitivity vector as calculated from the model predictions and the sensitivity vector of the experimental data. A low distance indicates that the coding properties have been reproduced well by the model.

### Cell Selection

In addition to excluding cells that could not be matched to one of the five analyzed cell types as noted above, we excluded cells from further analyses that responded unreliably during the saccadic stimulus or the full-field brightness steps. To measure a cell’s reliability during one of these stimuli, we first split its responses into odd and even trials and calculated PSTHs individually. In the case of the saccadic stimulus, we linked the 16 PSTHs of different starting and target position combinations together to generate a single PSTH for all odd trials and a single PSTH for all even trials. In the case of the full-field brightness steps, we only considered the 400 ms after any brightness change and linked the responses together to determine odd and even PSTHs in order to exclude long periods without changes in the stimulus. Next, we computed the coefficient of determination R^2^ between the odd PSTH as data and the even PSTH as prediction, and vice versa, and averaged these two values (Karamanlis and Gollisch, 2021). Any cell with an averaged R2 below 0.2 for either the saccadic stimulus or the full-field brightness steps was excluded from the analysis. In total, we analyzed 99 On parasol, 18 On midget, 113 Off parasol, 19 Off midget, and 9 Large Off cells from three experiments.

## Results

### Stimulus and Analysis

Saccades form rapid transitions between fixated images, and elicited responses in neurons of the visual system may be influenced by the pre-saccadic image, the post-saccadic image, and the transition in between. In order to investigate the coding properties of ganglion cells in the primate retina under saccade-like image transitions, we recorded ganglion cell spiking activity in isolated marmoset retinas with multielectrode arrays while projecting a saccade-like stimulus onto the photoreceptors. To systematically probe transitions between different illumination patterns inside the receptive fields of different ganglion cells, we chose a square-wave luminance grating with a bar width of 90 μm as the spatial layout of the stimulus. Taking into account the size of the marmoset eye (Troilo et al., 1993), this corresponds to approximately 0.9° visual angle or, for example, a 10 cm thick tree branch at a distance of about 6 m.

To mimic the sequential order of fixations and saccades, the grating remained stationary for a fixation period of 533 ms at one of four equally-spaced positions, which we call Position 1 to 4, before being shifted rapidly within 67 ms to a new position to start the next cycle of fixation and transition (Fig 1A). In half of the transitions, the shift itself was masked by a gray screen at the mean light intensity of the grating to probe for effects of visual stimulation during the transition. The order of fixation positions and the occurrence of the gray-screen mask were randomized. Altogether, there were 16 possible combinations of grating positions before (“starting position”) and after (“target position”) a transition, and each combination was presented with transitions by grating motion as well as by homogeneous illumination.

**Figure 1.**
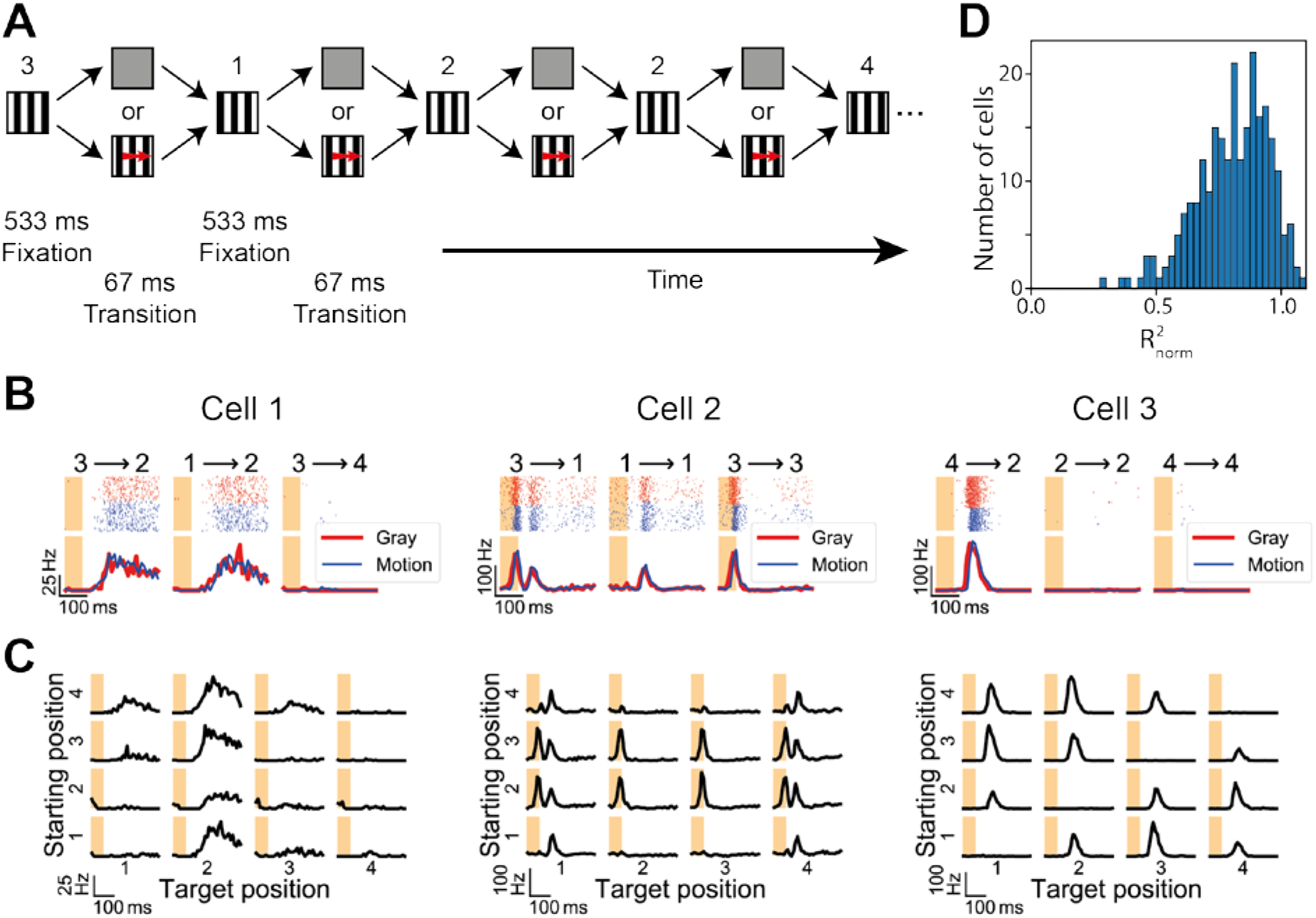
Sample ganglion cell responses to saccade-like grating shifts. (A) Schematic representation of the stimulus. The stimulus consisted of a sequence of fixations of a square-wave grating, which remained still for 533 ms at one of four possible positions (spatial phases), and brief transitions over 67 ms to the next fixation position. The transition either occurred via a rapid motion of the grating or via a gray-screen mask at mean luminance. The sequence of fixation positions and the type of transition were randomized. (B) Responses of three sample ganglion cells to different combinations of grating position before and after the transition. Top row shows raster plots for both gray (red) and motion (blue) transitions, bottom row shows the corresponding PSTHs. Shaded areas mark the transition periods. (C) PSTHs of the sample cells for all 16 possible combinations of grating positions before the transition (starting position) and after the transition (target position). Here, responses to gray-screen and motion transitions were pooled. (D) Similarity of responses to gray and motion transitions, assessed via the normalized coefficients of determination 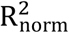 between responses to gray and motion transitions for all cells included in the analysis. 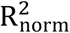 was calculated as the modified coefficient of determination between responses to gray and to motion, normalized by the modified coefficient of determination between the odd and even trials (independent of the transition type). Nearly all 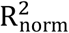 values are larger than 0.5, indicating high similarity between the responses to gray and motion transition.

We observed that ganglion cell responses could depend on both the grating position before and after the transition, but hardly depended on whether the transition occurred via motion or a gray screen. Some of the different response patterns and their dependencies on starting and target positions are exemplified by the three sample cells displayed in Fig 1B. Cell 1 responded strongly after transitions from Position 3 to 2 as well as from 1 to 2, but not for transitions from Position 3 to 4, suggesting a preference for a specific target position, namely here Position 2. When comparing firing-rate profiles for transitions via motion and via a gray screen, on the other hand, nearly identical responses were found for each individual combination of starting and target image. This was also the case for the two other sample cells of Fig 1 as well as for the entire population of ganglion cells (Fig 1D), except for a small latency effect in some cells (e.g. Cells 2 and 3), which played no role for our analysis of response strength and sensitivity to the pre- and post-saccadic images. We therefore made no further distinction between the two transition types and pooled responses from motion and gray-screen transitions.

In order to visualize the response characteristics more systematically, we computed the peri-stimulus time histograms (PSTHs) for all 16 combinations of starting and target position and displayed them in a matrix-like fashion (Fig 1C). In this depiction, it becomes immediately apparent that Cell 1 responded strongly when the target grating position was Position 2, but not for other targets, and that the response was only slightly modulated by the starting position. Thus, this cell is sensitive mostly to the post-saccadic image.

Other ganglion cells, like Cell 2, could display two distinct response peaks, one during the transition itself and one after the onset of the new fixation. For this cell, both peaks occurred when the grating switched from Position 3 to 1, but other sample transitions with a different starting or target position elicited only one or the other (Fig 1B, center). The matrix-like display of all 16 firing rate profiles (Fig 1C, center) reveals that the first peak was sensitive to the starting position, occurring systematically for Positions 2 and 3, whereas the second peak depended on the target position and was elicited by Positions 1 and 4. Note that due to the cyclical nature of the grating, Positions 1 and 4 are neighboring just like Positions 2 and 3 are.

For some ganglion cells, responses to the saccadic stimulus depended more intricately on the combination of pre- and post-saccadic images. Cell 3, for example, exhibited increased activity if the grating position changed from 4 to 2, but neither the starting nor the target position alone were sufficient to evoke a strong response if there was no change in the grating position across the transition (Fig 1B, right). Indeed, none of the starting or target positions by itself were sufficient to evoke a response of Cell 3, because starting and target position had to differ to trigger the cell (Fig 1C, right). Thus, this cell was sensitive to a change in the grating position, but invariant to the specific starting and target positions of the transition.

In order to systematically compare these response patterns for different types of ganglion cells, we sought a reduced, quantitative description which still captures the dependencies of the responses on the starting position, on the target position, and on whether there was a change of the position. To take the distinct early and late responses of some cells into account (e.g., Cell 2 in Fig 1), we analyzed the peak firing rates in two response windows, the first from 30 ms after transition onset to 30 ms (20 ms for one experiment) after the onset of the new fixation, and the second from 30 ms (20 ms) until 200 ms after the onset of the new fixation (Fig 2A).

**Figure 2.**
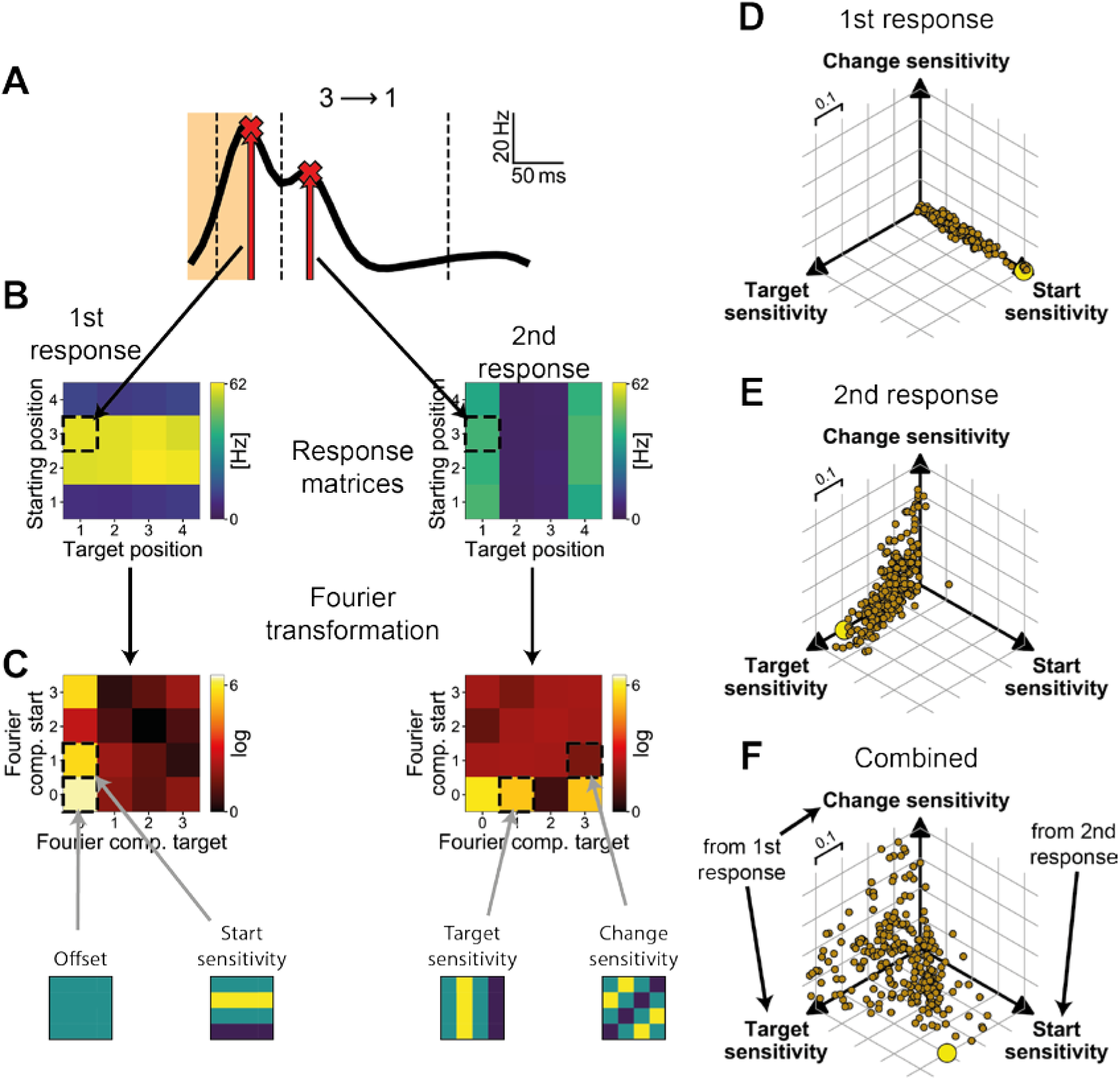
Analysis of ganglion cell sensitivity. (A) PSTH of a sample cell (Cell 2 from Fig 1) for the transition between Position 3 and Position 1, smoothed by a Gaussian filter. (In the quantitative analysis, the two response phases were smoothed separately.) Dashed vertical lines mark the boundaries of the first and second response phases. For each response phase, the peak response is identified as depicted by the red crosses and arrows. (B) The response matrices contain the peak response during the first phase (left) and second phase (right) for all combinations of starting and target position, depicted here in a color-coded fashion with brighter colors denoting stronger responses. The dashed squares indicate the entries that correspond to the sample PSTH shown in (A). (C) Fourier transformations of the response matrices. The entries of the transformed matrices quantify patterns in the response matrices. Entries that are highlighted by dashed squares correspond to relevant patterns in the response matrix, which are depicted schematically below the Fourier transformed matrices. For the sample cell, the yellow (0, 1) entry of the first and the yellow (1, 0) entry of the second Fourier transformed response matrix reflect the start and target sensitivity of the cell. (D) Scatterplot of the start, target, and change sensitivity of the first response for each cell. The large yellow data point marks the sample cell from (A-C). (E) Same for the sensitivity of the second response. (F) Elements of the final sensitivity vector for each cell, obtained by combining the start sensitivity of the first response and the target and change sensitivity of the second response.

To systematically analyze the dependence of the responses in these two temporal windows on the combination of starting and target position of the grating, we collected the peak firing rates of each window in a 4×4 matrix, corresponding to the 4×4 transitions from starting to target position (Fig 2B). The structure of these two response matrices contains information about the cell’s sensitivity to specific grating positions. For example, the first response matrix of Cell 2 from Fig 1, displayed in Fig 2B (left), contains horizontal stripes, demonstrating its sensitivity to specific starting positions (here Positions 2 and 3). The vertical stripes in the response matrix for the second time window (right), on the other hand, correspond to sensitivity to specific target positions.

The occurrence of stripes in the response matrices thus denotes the stimulus sensitivity during the selected response window. The specific position of the stripes, however, merely depends on the location of the cell’s receptive field relative to the bars of the grating. For example, Position 2 of the grating presumably brought an increase in preferred contrast to the receptive field of Cell 1 in Fig 1, but if that cell’s receptive field had been displaced by a quarter grating period in the right direction, the same response peak would have occurred for Position 3 instead.

Therefore, in order to make the analysis invariant to the receptive field position, we applied a two-dimensional Fourier transformation to the response matrices and only considered the amplitudes of the Fourier components. These capture the prevalence of stripe-like structures in the response matrices along the horizontal, vertical, or diagonal directions, while their phases, which contain the information about the positions of these structures, are discarded. The Fourier transformation yields another 4×4 matrix for each of the two time windows (Fig 2C). Three entries of the Fourier transformed response matrices are of particular interest as they correspond to the aforementioned sensitivities to starting position, target position, and change of position. Horizontal and vertical stripes in the response matrices, corresponding to sensitivity to the starting and target position, are captured by the (0, 1) and the (1, 0) entry, respectively, of the Fourier transform. For the sample cell of Fig 2, the former is correspondingly large for the first time window and the latter is large for the second time window. Note that, for symmetry reasons, the same values are found in the (0, 3) and (3, 0) entries. The entries (0, 2) and (2, 0), on the other hand, would correspond to higher harmonics in the sensitivity to starting and target position. It would also be possible that a response is sensitive both to a certain starting and a certain target position, which would correspond to large values for both the (0, 1) and the (1, 0) entry.

Cells sensitive to change of the grating position, like Cell 3 from Fig 1, have diagonal stripes in their second response matrix (data not shown; see inset at bottom right in Fig 2C for a schematic). This type of pattern is reflected in the (3, 1) entry of the Fourier transformed response matrix (as well as in the identical (1, 3) entry). In the case of Cell 3 of Fig 1, the large (3, 1) Fourier component reflects decreased activity along the diagonal of equal starting and target position. On the other hand, since our analysis disregards phase information of the Fourier transform, a large (3, 1) entry could also signify increased activity along this diagonal, corresponding to sensitivity to recurrence of the same grating position across the transition. Such sensitivity to image recurrence has indeed been described for certain ganglion cells of the mouse retina (Krishnamoorthy et al., 2017). For the present datasets from the marmoset, however, we did not find any image-recurrence-sensitive cells. A strong (3, 1) entry in our data always corresponded to a response sensitivity to a change of the grating position.

Note that the (1, 1) Fourier component would correspond to activity patterns along the other diagonal in the response matrix, from top-left to bottom-right. As expected, this component, as well as higher-harmonics, e.g., entries (2, 0) and (2, 1), rarely showed large values and did not provide information that helped characterize the responses. Thus, we found that focusing on the three Fourier components (0, 1), (1, 0), and (3, 1) was most effective. Finally, for some cells, none of the three entries described above contained a large value. In such a case, the cell either did not respond at all during the corresponding time window or responded indifferently to all transitions with no dependence on the starting or target position or any combination of the two.

To compare the sensitivities to pre-saccadic and post-saccadic images and combinations thereof across cells, we combined the three relevant entries of each Fourier transformed response matrix ((0, 1), (1, 0), and (3, 1)) into a three-component vector and normalized it (see Methods). For each response time window, its elements thus characterize a given cell’s sensitivity to the starting position, to the target position, and to change in the fixated stimulus pattern across the saccade. Examining this vector for the first response window for each ganglion cell (Fig 2D) shows that this response phase was generally only sensitive to the starting position, as the other two Fourier components for target position and change sensitivity were always near zero. This was expected, as the first response occurs too early to be affected by the new fixation and is thus mostly elicited by the offset of the starting position grating. The second response, on the other hand, was dominated by the components corresponding to sensitivity to the target position and to change, with considerable differences in the magnitude of these two components between individual cells, but with generally little sensitivity to the starting position alone (Fig 2E). Thus, this response component is typically affected by the target position of the grating and by combinations of the starting with the target position, consistent with its occurrence several tens of milliseconds after the onset of the new fixation.

To jointly analyze the most relevant patterns of stimulus sensitivities during the first and second response window, we thus combined the sensitivity for the starting position of the first response with the target and change sensitivities of the second response to obtain a final three-component vector, which we call the sensitivity vector. The sensitivity vector describes the most pronounced response properties of a ganglion cell under our saccade stimulus (Fig 2F). For example, the sensitivity vector for the sample cell of Fig 2A-C (yellow dot in Fig 2D-F) has large sensitivity values for the starting and the target position, but not for the change of position, reflecting the horizontal and vertical stripes in the response matrices of Fig 2B and the lack of a diagonal structure.

### Responses of Different Cell Classes

We next asked whether the observed differences in response sensitivities were connected to the different types of ganglion cells. To investigate this question, we first classified cells according to standard response characteristics measured under spatiotemporal white-noise stimulation. Specifically, we measured the cells’ spatial receptive field and temporal filter via the spatiotemporal spike-triggered average, their output nonlinearities, and their spike autocorrelations (Fig 3, see Methods for details). Five distinct classes could be readily identified, including the standard types On and Off parasol as well as midget cells. On and Off parasol cells displayed fast, biphasic temporal filters and receptive field tiling. On and Off midget cells had slower filters and smaller receptive fields. Here, however, tiling was not apparent, owing to the limited number of recorded cells. In addition to these four major primate ganglion cell types, we also identified a fifth type, an Off cell with slow temporal filters and large receptive fields. We here refer to this type as Large Off cells.

**Figure 3.**
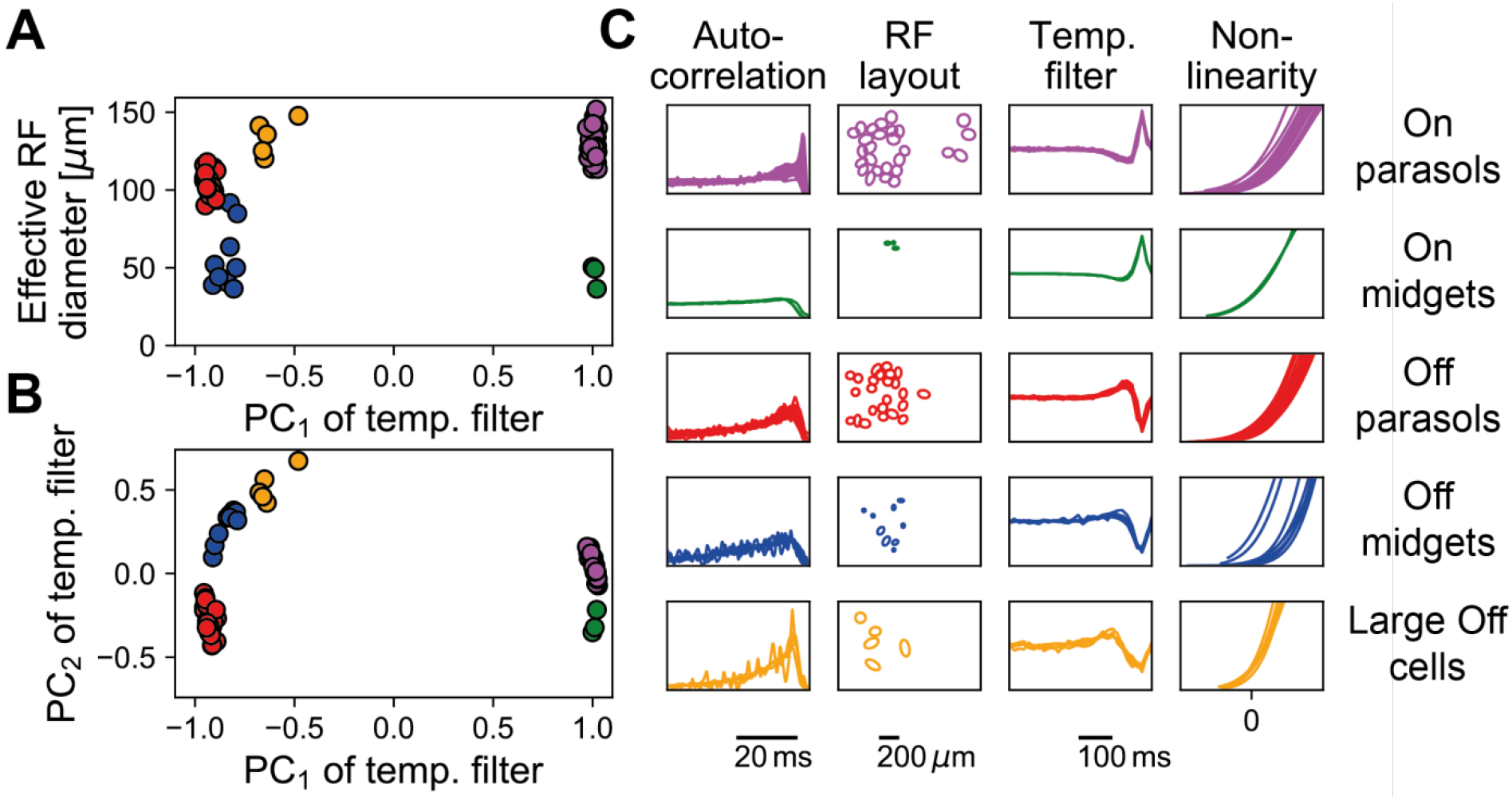
Classification of retinal ganglion cells of a sample experiment. (A) Scatterplot of the effective receptive-field diameter versus the projection onto the first principal component of the temporal filter for all classified cells (magenta: On parasol cells, green: On midget cells, red: Off parasol cells, blue: Off midget cells, orange: Large Off cells). (B) Same as (A), but for second principal component versus first principal component of the temporal filter. (C) Autocorrelation functions, receptive field layouts (1.5-sigma ellipses of receptive-field Gaussians), temporal filters, and nonlinearities of all classified cells, grouped by cell type.

We found that the identified major cell types exhibited distinct, characteristic responses to the saccadic stimulus. Figure 4 shows representative response profiles, which illustrate the differences in response patterns between the cell types. Many On parasol cells had two separate response components, a first response sensitive to the starting position and a second response sensitive to the target position. For example, the first On parasol cell in Fig 4A (left column, top) displayed an early response peak if the starting position was 4 (weaker if it was 1 or 3) and a later response peak if the target position was 2. Other On parasol cells did not show a clear preference for specific starting or target positions and instead responded rather indifferently (Fig 4A, left column, bottom example). The sensitivity vectors of all On parasol cells show that there was a continuum between these two response types with indifferent cells lying closer to the origin (Fig 4B, left column). This range in response characteristics seems to be related to variations in receptive field size of On parasol cells across our recordings. In fact, there was a pronounced correlation between receptive field size and the starting as well as target sensitivity of the cells (Fig 5A, B): larger cells usually responded more indifferently. This makes intuitive sense, as larger receptive fields are more likely to contain both dark and bright bars of the grating for each position.

**Figure 4.**
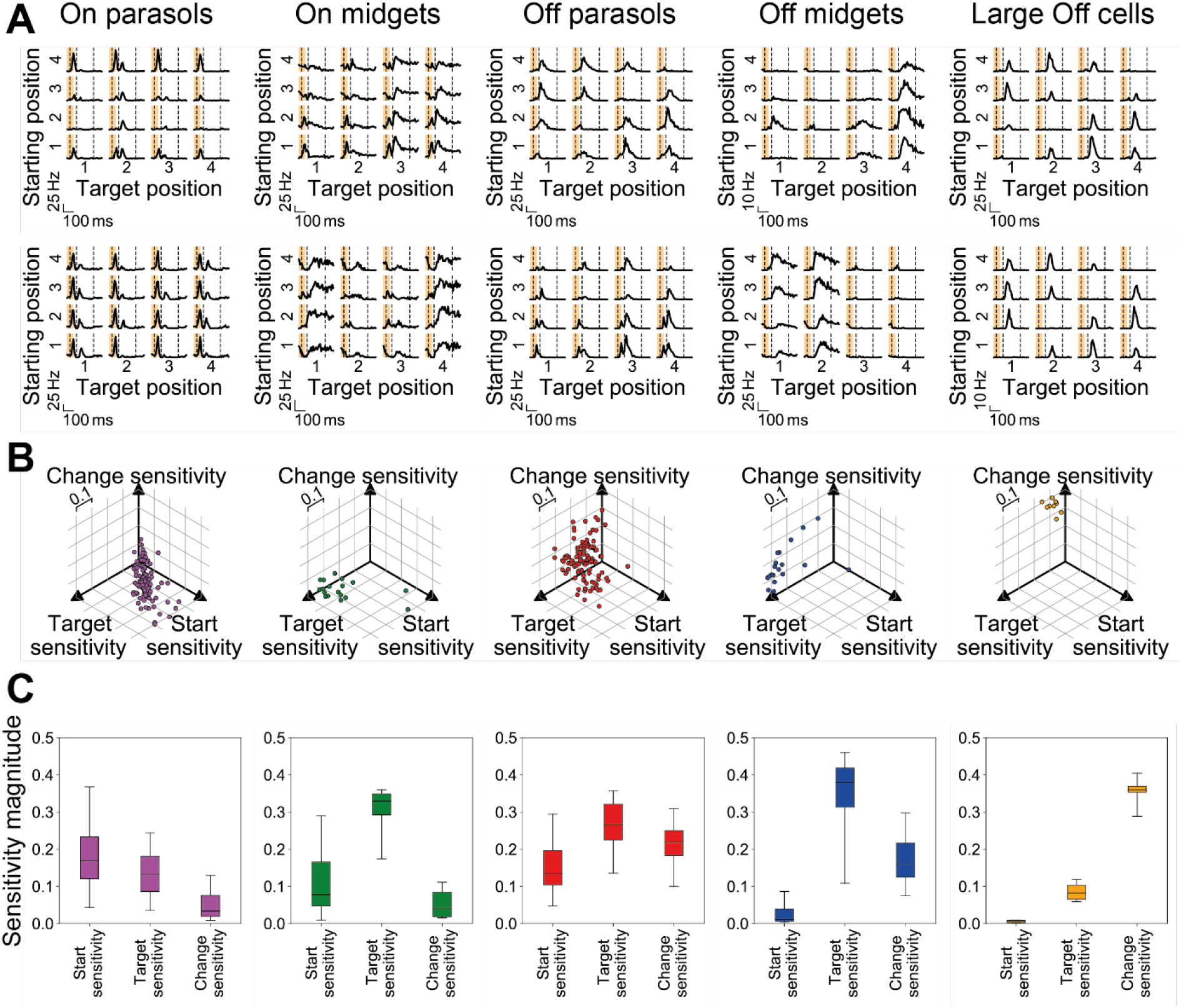
Coding properties of ganglion cell types. (A) PSTHs of different sample cells, showing the responses of the cells to all 16 combinations of starting and target position. Shaded regions denote the transition period and dashed vertical lines mark the borders of the response phases used for analysis. (B) Sensitivity vectors of all cells of the five distinguished cell types. (C) Boxplots of the distributions of sensitivity measures (entries of the sensitivity vector) for each of the five cell types. Boxes denote the central 50% of data points (i.e., from 25% to 75%), whiskers the central 90% (from 5% to 95%), and horizontal lines inside the boxes the medians.

**Figure 5.**
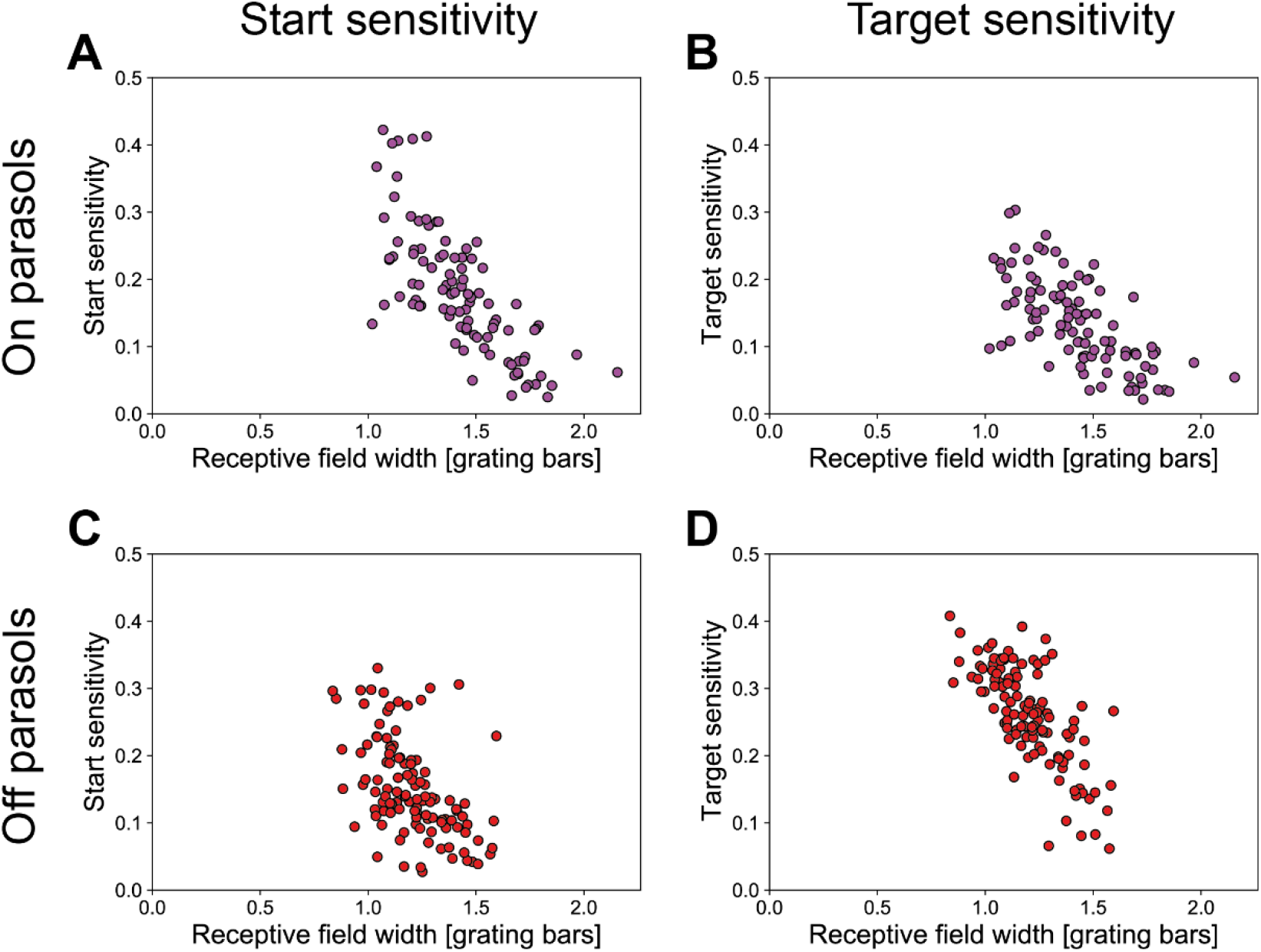
Dependence of sensitivity on receptive field size. (A) Dependence of the sensitivity to the starting position on the width of the receptive field for all On parasol cells. Receptive field width was defined as the extent in x-direction (perpendicular to the grating) of the 1.5-sigma ellipse of the receptive-field Gaussian and given relative to the size of a grating bar.(B) Same as in (A) but for sensitivity to the target position. (C) same as in (A) but for Off parasol cells. (D) Same as in (A) but for Off parasol cells and sensitivity to the target position.

Fig 4C (left column) shows boxplots of the sensitivities of On parasols confirming that they are mostly sensitive to the starting position and (somewhat less) to the target position. Change sensitivity only plays a subordinate role. In one experiment, though, we also found some On parasol cells whose second response did not seem to be sensitive to the target position but rather to the change of position (data not shown).

Similar to On parasol cells, On midget cells (Fig 4A, second column) also showed a first response sensitive to the starting position and a second response sensitive to the target position. In contrast to their parasol counterparts, however, the responses of On midget cells were dominated by the second response peak, which was more pronounced and sustained than the first. This shifts the sensitivity balance towards the target position (Fig 4B, C).

For Off parasol cells, the most striking response feature was that many cells were sensitive to a change of the grating position across the transition. This is evident from the reduced responses during the second response window on the diagonal of equal starting and target position in the matrix representation of the PSTHs (Fig 4A, third column) as well as from the large change sensitivity component of the sensitivity vectors (Fig 4B, C). In addition, however, there often was also considerable sensitivity to the specific starting and target positions in the first and second response, respectively. Which of these two response characteristics was the more prominent depended on the receptive field size (Fig 5C, D). Cells with larger receptive fields were usually dominated by their strong sensitivity to the change of the grating position while the specific starting and target positions did not significantly influence the responses (Fig 4A, top example). Smaller Off parasol cells, however, were also sensitive to the actual starting and target positions, which obscures the change sensitivity to some degree (Fig 4A, bottom example). Again, the dependence on receptive field size makes intuitive sense, as the activation of larger receptive fields is less dependent on the exact grating position.

Off midget cells were mainly sensitive to the target position. Both examples in Fig 4A (fourth column) show cells that responded only to the occurrence of one or two specific target positions with only some modulation by the starting position. The moderate amount of modulation by the starting position was such that responses to a recurrence of the same grating position were reduced. This mild change sensitivity is also revealed by the sensitivity vectors (Fig 4B, C). Overall, however, Off midget cell responses were dominated by their target sensitivity, which they displayed more strongly than any of the other cell types.

For the Large Off cells, the striking feature was their pronounced sensitivity to the change of the grating position (Fig 4A, right column) with essentially no sensitivity to the specific starting or target position (Fig 4B, C). These cells generally showed no activity during the first response window and transient responses during the second whenever starting and target position differed.

Together, these analyses show that different types of ganglion cells systematically differ in how the combination of pre- and post-saccadic images affect the spiking activity during and after a saccade-like image transition.

### Modeling Responses to the Saccadic Stimulus

We next asked to what degree the different response patterns of the distinguished cell types followed from how the cells responded to simple steps in light intensity. This might yield hints to circuit structures that are triggered specifically by saccades. As the saccade-like motion transition of the stimulus evoked essentially the same responses as the gray transition (Fig 1), the saccadic stimulus can be treated as a combination of an offset of bright and dark regions followed by an onset of a new bright/dark pattern. We therefore compared the responses to the saccadic stimulus with responses to full-field brightness steps from mean light intensity (gray) to high intensity (white) or to low intensity (black) and back to gray. As expected, On cells responded to an increase in brightness whereas Off cells responded to a decrease and parasol cells responded more transiently than midget cells (Fig 6A). The Large Off cells also responded transiently to decreases of the brightness, but with a longer latency than Off parasol cells.

**Figure 6.**
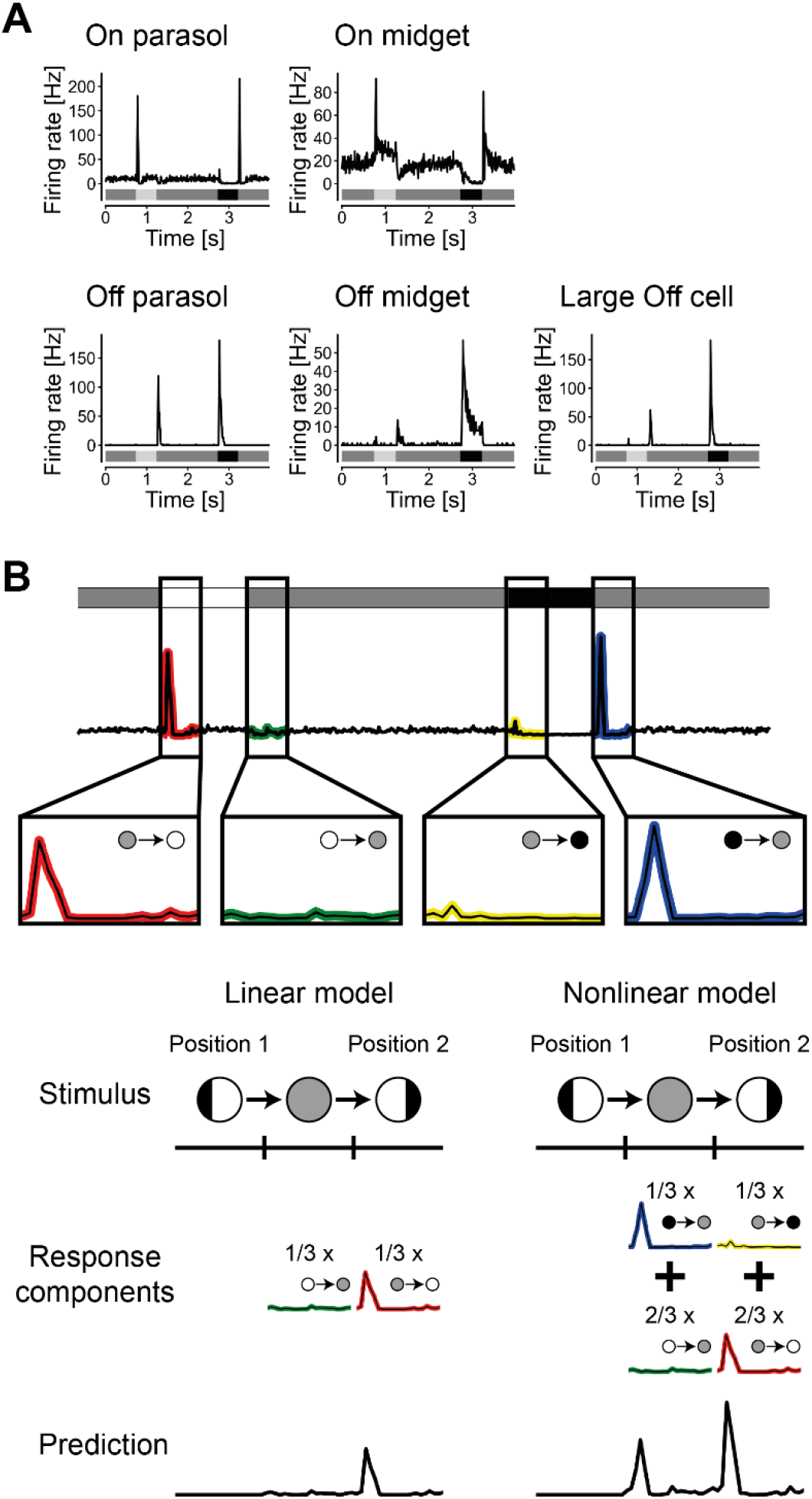
Modeling ganglion cell responses to saccades based on responses to brightness steps. (A) Exemplary PSTHs for all five retinal ganglion cell types to full-field brightness steps. The stimulus is schematically depicted beneath each PSTH. (B) Schematic depiction of obtaining a response prediction for the ON parasol cell in (A). The cell’s response to the full-field brightness steps (top) was split into responses to the on- and offsets of white and black (zoomed-in insets below, small circles and coloring of the PSTHs denote the brightness change). For the linear model (left), the response to an offset of the first grating was estimated by scaling the cell’s response to the offset of the appropriate brightness step, here one third of the offset of white, corresponding to the relative decrease in mean luminance inside the receptive field. Analogously, the response to the onset of the new grating was estimated here as one third of the onset of white, corresponding to the increase in mean brightness. Unlike depicted here, these two response components overlapped strongly, because of the briefness of the transition. They were then summed to form the final response prediction. For the nonlinear model (right), each pixel directly contributed response components to the final response thereby omitting the averaging of the brightness inside the receptive field. At the offset of the first grating in this example, one third of the receptive field turned from black to gray and two thirds from white to gray, yielding a one-third contribution of the black offset response and a two-thirds contribution of the white offset response. The onset of the second grating position was treated analogously and the four response components were summed to generate the final response prediction.

To assess the relation between the responses to brightness steps and responses to the saccadic stimulus, we aimed at modeling the latter based on the former (Fig 6B). Since the saccadic stimulus contains spatially structured images and given that ganglion cells can pool signals over space either linearly or nonlinearly (Enroth-Cugell and Robson, 1966; Hochstein and Shapley, 1976; Schwartz and Rieke, 2011; Gollisch, 2013; Turner and Rieke, 2016; Karamanlis and Gollisch, 2021), we correspondingly set up two models with either linear or nonlinear spatial integration. The linear model employed a linear receptive field, which was obtained by separating the spike-triggered average (STA) under spatiotemporal white-noise stimulation (Chichilnisky, 2001) into a spatial and a temporal filter and fitting a two-dimensional Gaussian to the spatial filter. The grating stimulus was then filtered by the receptive field to obtain an effective average brightness of the pre-as well as of the post-saccadic grating inside the receptive field. This means that bright and dark regions inside the receptive field can partially cancel each other out. The filtering yielded a sequence of step-like brightness changes, and the cell’s response was modeled by combining the measured responses to the corresponding brightness steps, scaled according to the effective average brightness of the grating. The second model employed a nonlinear receptive field. Here, each pixel within the receptive field contributed individually to the firing rate according to the brightness changes that occurred at that pixel, without any cancelation by other pixels. The contributions from all pixels were then summed in a weighted fashion according to the receptive field of the cell. Thus, the linear model predicts responses only according to the overall brightness changes in the receptive field, whereas the nonlinear model also includes the spatial structure of a stimulus in its response.

We found that most of the responses could be predicted by at least one of the two models. The responses of On parasol cells were captured well by the nonlinear model, but not by the linear model. The linear model failed, for example, to recreate the strong responses of the sample On parasol of Fig 7A (left) to transitions between Positions 2 and 4 (Fig 7B). These grating positions yielded roughly equal bright and dark contrast in the cell’s receptive field (Fig 7D), and the linear model therefore predicted no response, because these regions can cancel each other out. By contrast, the nonlinear model correctly captured the response patterns (Fig 7C). It also succeeded in predicting a first response sensitive to the starting position and a second response sensitive to the target position, even though it underestimated the strength of modulation of the second response caused by the target position.

**Figure 7.**
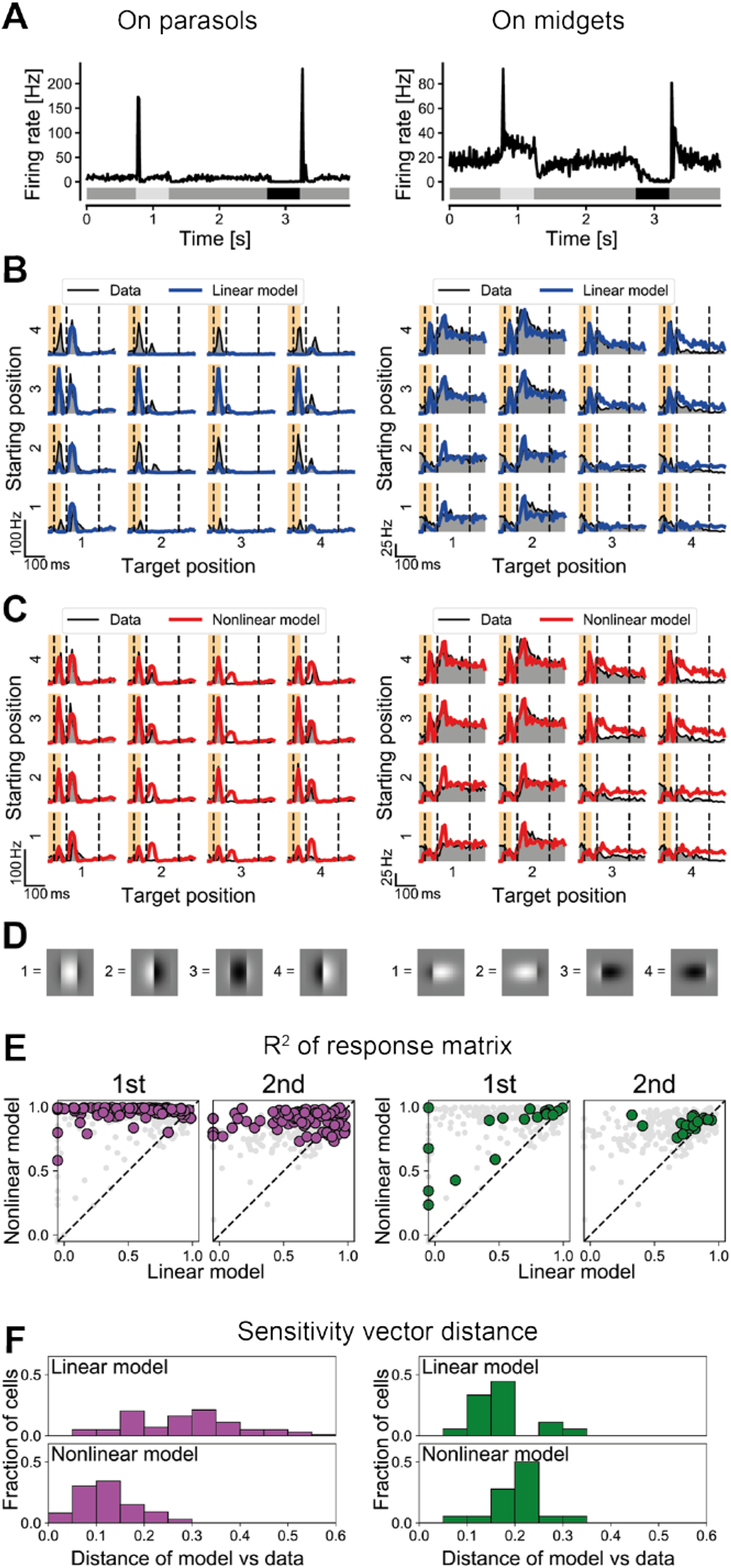
Model evaluation for On parasol (left) and On midget (right) cells. (A) PSTHs of sample cells to the full-field brightness steps. (B) Experimental responses to the saccadic stimulus (thin black line with gray filling) and predictions by the linear model (thick blue line) of the same sample cells to the saccadic stimulus. Layout of the plot is the same as in Fig 1C. (C) Same as (B) but for the nonlinear model (red line). (D) Illustrations of the contents of the cells’ receptive fields at the four grating positions. (E) Modified coefficient of determination R^2^ for the response matrix of the data versus the linear model (x axis) and versus the nonlinear model (y axis). R2 values are shown separately for the first and second response matrix. Colored dots represent all ganglion cells of the column’s cell type, light gray dots all other cell types. Cells with a model R2 below −0.05 have been plotted on the axis. (F) Distributions of the distances between modeled and experimental sensitivity vectors for the linear and the nonlinear model.

In order to quantify the accuracy of the model predictions, we computed modified coefficients of determination R^2^ between the response matrices of the experimental data of a cell and the response matrices as calculated from the modeled PSTHs (see Methods). For each response phase, this yielded one value per cell and model, which usually lay in the range of zero (no correlation between model and data in that response phase) to unity (perfect correlation between model and data). For the On parasol cells, the nonlinear model, but not the linear model, generally achieved R2 values close to unity, especially for the first response phase and only slightly less so for the second response phase (Fig 7E), corroborating the nonlinearity of On parasol receptive fields under these stimulus conditions.

While the computed coefficients of determination quantify how well the amplitudes of the response peaks in the PSTHs are captured, they do not directly assess whether the models capture a cell’s sensitivity characteristics with respect to starting position, target position, and change of the grating. As an alternative measure of model accuracy, we therefore computed the distance between the measured sensitivity vector of a cell and the sensitivity vector calculated from the modeled responses. A small distance indicates that the response sensitivities have been reproduced, while a large distance represents discrepant sensitivities. For the On parasol cells, the distance of the linear model’s sensitivity vectors to the experimental sensitivity vectors was generally large, while the nonlinear model produced small distances (Fig 7F). This confirms that On parasol responses to the saccadic stimulus could be modeled well by using the responses to a full-field stimulus and assuming a nonlinear receptive field. Only in one experiment, some On parasol cells with unusually slow and weak responses to the full-field brightness step from black to gray were not modeled well by the nonlinear model (data not shown).

On midget cells, like the example in Fig 7A-C (right), were modeled decently by both the linear as well as the nonlinear model. Here, the linear and the nonlinear model yielded similar response predictions because the small receptive fields of these cells contained mostly only a single bar of the grating (Fig 7D). Due to the lack of spatial structure within the receptive field, the nonlinearity played hardly any role. While the strength of the sample cell’s second response was partially overestimated, the main response properties, i.e. the transient, start-sensitive first response and the sustained, target-sensitive second response, were successfully predicted. For both response phases, the models achieved relatively high R2 measures of the response matrices, with a slight tendency towards better predictions by the nonlinear model (Fig 7E; p=1.5×10^-5^ and p=3.3×10^-4^ for first and second phase, respectively, Wilcoxon signed-rank test). The distance of sensitivity vectors, on the other hand, was smaller for the linear model (Fig 7F; p=1.9×10^-3^, Wilcoxon signed-rank test).

In contrast to On cells, the responses of Off parasol as well as Large Off cells could only partially be explained by the linear or nonlinear model. For Off parasol cells and Large Off cells, the linear model displayed similar problems as for On parasol cells, often strongly underestimating the responses (e.g. Fig 8B) due to cancelation that does not occur in the nonlinear receptive fields of these cells. By contrast, the nonlinear model could mostly reproduce the sensitivities to the starting and target position (Fig 8C). However, the responses on the diagonal, i.e., to a recurrence of the grating position, were consistently overestimated. For example, the Off parasol cell of Fig 8 lacked a strong second response to the transition when starting and target position were both Position 4. Both models predicted such a response, since the receptive field returned to being mostly filled with black after the transition (Fig 8D). Accordingly, both the similarity between response matrices (Fig 8E) and the sensitivity vector distance (Fig 8F) show that the models did not capture the response characteristics of Off parasol and Large Off cells as successfully as for other cell types.

**Figure 8.**
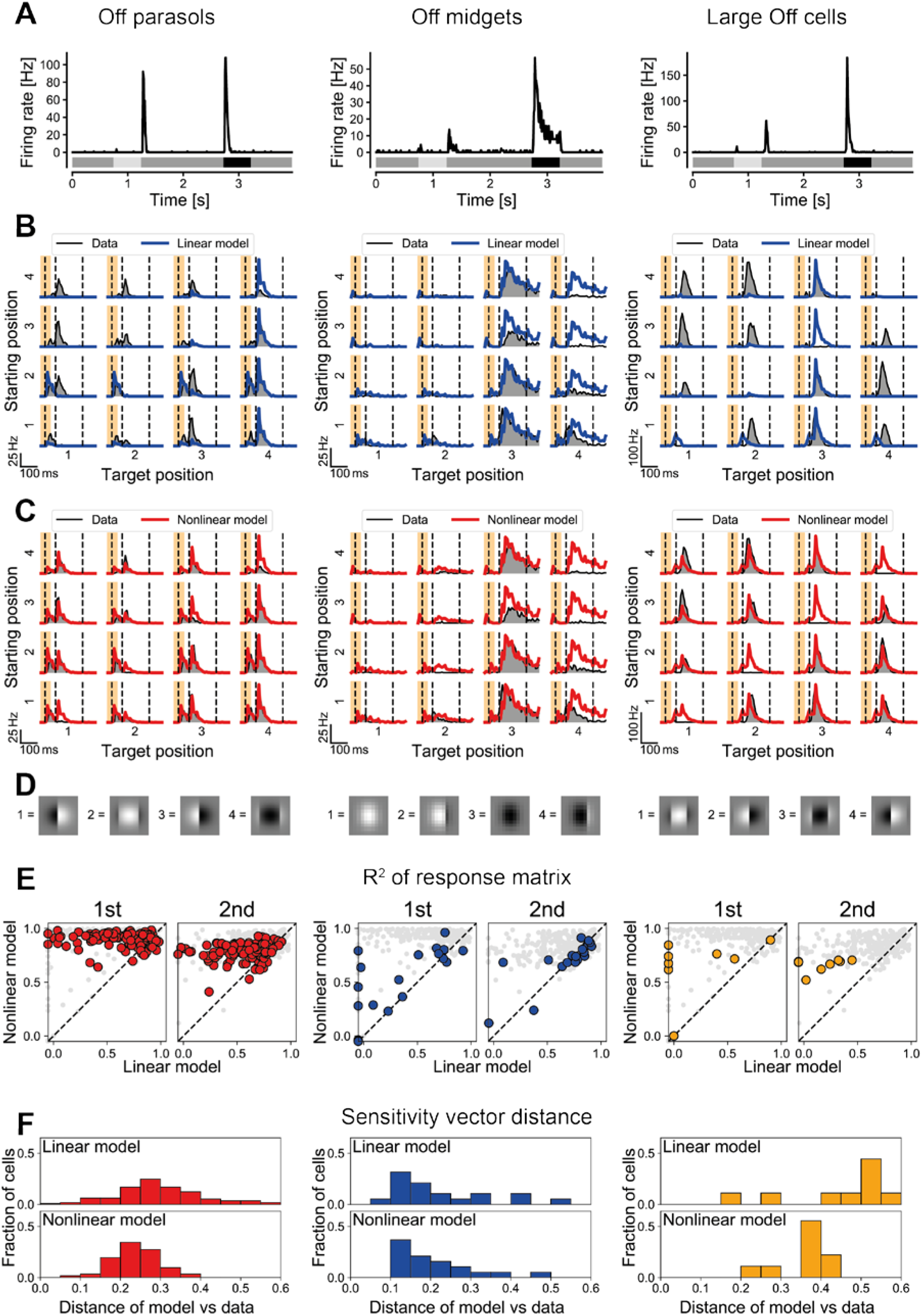
Model evaluation for Off cells. Same layout and subfigures as in Fig 7, but with Off parasol cells in the left column, Off midget cells in the middle column, and Large Off cells in the right column.

For Off midget cells, akin to On midget cells, the small size of their receptive fields led to similar predictions by the linear and nonlinear model (Fig 8A-D, middle column). For the sample Off midget cell, the general target sensitivity was reproduced, although responses to Position 4 as target position were overestimated, possibly a result of noise in the receptive-field measurement. Furthermore, the slight modulation of the responses by the starting position hinting at some change sensitivity was not captured by the models. For the first response phase, the models achieved comparatively low R2 values (Fig 8E), largely because Off midget cells responded only weakly and unreliably during this phase. For the second response phase, however, which included the bulk of the Off midget responses, both models achieved decent R2 values, but were likely suffering somewhat from the mild change sensitivity in the responses that was not captured by the models. The distance between the modeled and measured sensitivity vectors of the Off midget population was rather small (Fig 8F), indicating that the general sensitivity profiles of Off midget cells could mostly be explained by their responses to the full-field brightness steps.

Figure 9A-C summarizes the sensitivity measures predicted by the two models and directly extracted from the data. Evidently, while the sensitivity to starting and target position could generally be explained by a cell’s preference for light increments and decrements (Fig 9A, B), in particular when using the nonlinear model, the change sensitivity could not (Fig 9C). Accordingly, the sensitivity vectors that were calculated from the model responses were largely restricted to the plane spanned by starting and target sensitivity; the predicted change sensitivity was always close to zero (Fig 9D, E). Therefore, the measured change sensitivity of Off parasol and Large Off cells and to a lesser degree also of Off midget cells appears to be the result of additional mechanisms. These mechanisms have the effect of spreading out the sensitivity vectors in the analyzed three-dimensional sensitivity space (Fig. 9F), thus diversifying the response characteristics of the different cell types under saccade-like image shifts.

**Figure 9.**
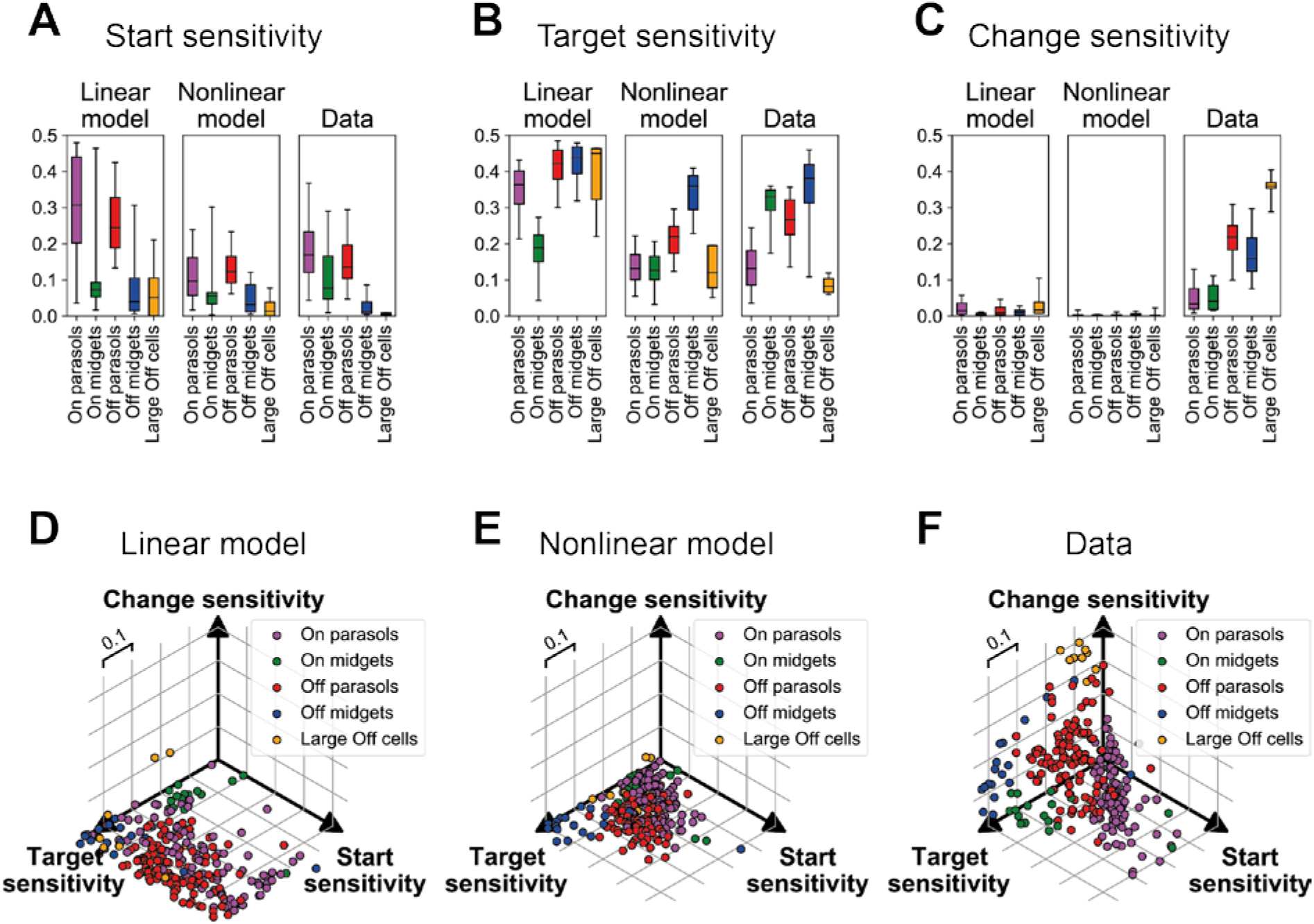
Sensitivity vectors from models and experiments. (A) Boxplots for the distribution of start-sensitivity values for each cell type, obtained via each of the two models and from the experimental data. The box marks the central 50%, whiskers the central 90%, and the horizontal line inside the box the median. (B) Same as (A) but for target sensitivity. (C) Same as (A) but for sensitivity to grating position change. (D) Scatter plot of sensitivity vectors calculated from the linear model for all cells of the five types. (E) Same as (D) but for nonlinear models. (F) Same as (D) but for experimental data.

## Discussion

Saccades pose a unique challenge to the visual system by presenting a rapid transition between two fixated images, separated by only about a hundred milliseconds or less. Despite their ubiquity in nearly all visual animals (Land, 1999), surprisingly little is known about how neurons in the visual system combine information from the pre- and post-saccadic images and what aspects of the two images are encoded in their responses in this context. In the present work, we have shown that responses of ganglion cells in the marmoset retina under saccadic stimulation do not simply represent the new fixation, but display a range of different dependencies on both the pre- and post-saccadic image (Fig 1). Quantifying the sensitivity to the starting position, the target position, and change across the transition (Fig 2) revealed that different ganglion cell types systematically displayed different sensitivity patterns (Fig 3–5). Using simple models with linear and nonlinear stimulus integration over space (Fig 6), showed that the dominant sensitivities of parasol and midget On cells could be reproduced based on the cell’s responses to isolated flashes of light intensity (Fig 7). By contrast, for many Off cells, especially Off parasol and a class of Large Off cells, the models failed to account for the observed sensitivity to change across the transition (Fig 8). Thus, the Off cells’ change sensitivity appears to require more complex circuit mechanisms, which are not triggered under isolated light-intensity flashes. This entails a new asymmetry in the functional properties of On and Off ganglion cell classes and contributes to diversifying the response patterns to saccade-like image transitions (Fig 9).

### Retinal Coding of Image Shifts

In search of the origins of saccadic suppression, multiple studies in various non-primate vertebrates have looked at the influence of saccades on the response strength of ganglion cells and found a diverse picture of enhancement, suppression, and indifference (Roska and Werblin, 2003; Amthor et al., 2005; Sivyer et al., 2019; Idrees et al., 2020). Fewer studies have investigated what ganglion cells encode during or after saccades, despite the likely importance of fixation onset for eliciting informative responses (Segev et al., 2007). Moreover, saccade-like image shifts may alter the message conveyed by ganglion cell spikes, as observed in the salamander retina, where On-Off ganglion cells were found to transiently switch their relative sensitivity to On-type versus Off-type stimuli after an image shift (Geffen et al., 2007). In an early study in the cat, Noda and Adey (1974) found sustained cells (probably X cells) that responded to preferred contrast in the target image, and transient cells (probably Y cells) that signaled the occurrence of a saccade. This is reminiscent of our findings of Off midget cells dominated by their target sensitivity and On parasol cells with large receptive fields that responded indifferently.

In a previous study from our lab, we had identified ganglion cells in the mouse retina that responded distinctly to the recurrence of an image (Krishnamoorthy et al., 2017). In the present study of the marmoset retina, we did not find such image-recurrence-sensitive (IRS) cells. In mice, IRS cells seem to correspond to transient Off alpha cells (Krishnamoorthy et al., 2017), which share some parallels with the Off parasol cells of the primate retina (Crook et al., 2008b). Here, however, we observed that Off parasol responses are sensitive to a change of the image, not to a recurrence. This corroborates the divergence between mouse and primate ganglion cell types (Peng et al., 2019), which may reflect the different visual requirements of these species (Baden et al., 2020).

### Large Off Cells

In addition to the standard midget and parasol ganglion cells, we also identified a fifth cell type, which we called Large Off cells. We distinguished these cells from Off parasol cells because they had slower temporal filters, larger receptive fields, and did not match Off parasol tiling. The identity of these cells is unknown, but the similarity of their response to Off parasol cells could suggest that they might be Off smooth monostratified (Off SM) ganglion cells, though other candidates, e.g., Off narrow thorny ganglion cells, also exist (Dacey, 2004; Masri et al., 2019; Grünert and Martin, 2021). In the macaque retina, Off SM cells have been described as similar to Off parasol cells, but with a longer latency and larger receptive fields (Crook et al., 2008a). In addition, SM cells tend to have irregular receptive fields with a hotspot structure (Rhoades et al., 2019), matching our observation that the Large Off cells in our recordings had more irregular receptive fields than parasol cells (data not shown). Yet, we did not find the counterpart of the Off SM cells, the On SM cells, and the difference in receptive field size between Large Off and Off parasol cells was smaller than what would have been expected from Off SM cells in the macaque retina (Crook et al., 2008a; Rhoades et al., 2019). The latter, however, might be a species-specific difference between macaque and marmoset or might result from differences in retinal eccentricity.

### Potential Mechanisms

The model analysis showed that the sensitivity to the starting and target position could largely be explained by a cells’ responses to full-field brightness steps, at least when nonlinear spatial integration is accounted for. Some differences between the cell types here are due to the speed of the responses. The fast response kinetics of the On parasol cells, for example, allow for a strong and distinct response to the onset of the transition with corresponding pronounced starting sensitivity, whereas the slower Off midget cells respond mostly only after the new fixation has started and are thus more sensitive to the target position.

Less clear is what the mechanism behind the change sensitivity observed in Off cells, in particular Off parasol and Large Off cells, might be. One hypothesis could be that transitions with no net change in the image pattern are simply too brief to be detected by the temporal filters of the ganglion cells, in particular since we did not observe different responses for motion and gray transitions. However, the temporal filters that we extracted from the STA typically peaked well before 67 ms, which is the duration of the transition. In addition, distinct responses to the onset as well as the offset of the transition are visible for most cell types. Therefore, it seems unlikely that the temporal filters are so slow as to cause change sensitivity.

Alternatively, neuronal or synaptic fatigue of local excitatory inputs accrued during the fixation of several hundred milliseconds prior to the transition might prevent responses to the new fixation when the same image recurs. However, the pronounced transiency of responses in Off parasol and Large Off cells and the lack of sustained activity speak against strong presynaptic activity that could trigger the required fatigue, which would need to be strong enough, for example, to prevent any response to recurring grating positions in Large Off cells.

Instead, we hypothesize that change sensitivity is caused by inhibition and propose a mechanism of local delayed crossover inhibition. In this mechanism, the Off bipolar cells that provide excitatory input to the change-sensitive Off cells receive inhibitory input from slow, narrow-field On-type amacrine cells. For recurring grating positions, this means that the local excitation from the dark stripes of the grating at the onset of the new fixation will be suppressed by inhibition that was triggered by the brightening at the same locations when the pre-saccadic grating disappeared. This previously triggered inhibition will not yet have decayed, if the activity of the corresponding amacrine cell is sustained enough to last across the duration of the transition.

Such crossover inhibition is the dominant inhibitory input to parasol cells and serves various functions by shaping ganglion cell responses in many species (Manookin et al., 2008; Werblin, 2010; Crook et al., 2011; Cafaro and Rieke, 2013; Rosa et al., 2016). Our hypothesized mechanism employs crossover inhibition onto bipolar cells as observed, e.g., in the rabbit retina (Wässle and Boycott, 1991; Molnar and Werblin, 2007), and act locally within receptive field subunits. This makes glycinergic narrow-field amacrine cells with their comparatively small receptive fields (Pourcho and Goebel, 1985; Menger et al., 1998; Masland, 2012) the likely candidate source. Known examples of narrow-field amacrine cells that implement crossover inhibition to Off bipolar cells exist. For instance, as shown in cat, rabbit, and rat retina, the AII amacrine cell receives On input from bipolar cells and provides glycinergic inhibition to Off bipolar axon terminals (Kolb and Famiglietti, 1974; Demb and Singer, 2012). A similar circuitry is also present in the macaque retina (Wässle et al., 1995).

### Limitations and Future Directions

The stimulus used here differs from real saccades in two important aspects. Firstly, the transition is not a real motion stimulus due to the limited framerate of our projection system. Yet, the high speed of a saccade and the corresponding motion blur make it likely that true saccadic motion and homogeneous illumination at mean light level are nearly equivalent stimuli for the retina, and we therefore do not expect this to strongly influence the findings regarding the encoding of pre-and post-saccadic images. Secondly, the applied gratings are artificial patterns, whose activation of the retinal circuitry may differ from that of natural stimuli (Turner and Rieke, 2016; Yu et al., 2022), though analyses of macaque ganglion cell responses to natural scenes containing self-motion signals found good correspondence of response characteristics with those typically obtained with simpler, artificial stimuli (Schottdorf and Lee, 2021). For the present work, the periodic nature of the gratings proved useful in allowing us to apply Fourier analysis for systematically analyzing the sensitivity profile of each cell independent of the particular position of a cell’s receptive field. This may pave the way for future investigations of responses to saccades with natural images.

Other future variations of the stimulus could help elucidate the circuit mechanism behind the observed change sensitivity. In particular, including gratings with a smaller spatial frequency would allow probing the small midget cells with stimuli that contain substantial spatial structure in the receptive field and elucidate whether Off midget cells are similarly change-sensitive as Off parasol and the Large Off cells. Furthermore, probing longer and shorter transition periods could reveal the timescale of the change sensitivity and relate this to potential inhibitory mechanisms. Finally, to test if crossover inhibition is involved in generating the change sensitive responses of Off cells, blocking the On pathway with L-AP4 (Slaughter and Miller, 1981) could be applied to remove On-type inhibition, even though this would not reveal specifics of the potential crossover circuitry.

### Asymmetry of the On and Off Pathways

The On and Off pathways in the retina had originally been viewed as symmetric, i.e., exhibiting the same response properties but for light increments and decrements, respectively (Schiller, 1992). Later, however, several studies found asymmetries between the On and Off pathways, such as differences in the spatiotemporal receptive field properties of macaque On and Off parasol ganglion cells (Chichilnisky and Kalmar, 2002). These differences might be linked to different connectivity in the underlying circuitry of the On and Off pathways found in primates and other species (Molnar and Werblin, 2007; Khuc-Trong and Rieke, 2008) and to the more strongly rectified synaptic input received by primate Off parasol cells, but also, e.g., guinea pig Off Y-cells, compared to their On counterparts (Zaghloul et al., 2003; Turner and Rieke, 2016). Importantly, these asymmetries extend to relevant functional differences like the encoding of natural images (Turner and Rieke, 2016).

While asymmetries between On and Off parasol cells (or Y-cells in other species) have been described previously, the midget pathways have received less attention. Here, we found asymmetries between the On and Off pathways of both parasol as well as midget ganglion cells. While Off midget cells were strongly sensitive to the target image with some change sensitivity, On midget cells were not change sensitive, but responded transiently to the preferred starting image. The asymmetric responses of parasol cells were even more striking. While On parasol responses represented images before and after a transition successively, Off parasol cells performed a computation across the transition by responding specifically to a change of the image. The different response characteristics suggest that On and Off cells encode different features of the visual stimulus in the context of saccades, similar to recent suggestions about the functional benefit of differences in spatial integration between On and Off parasol cells (Yu et al., 2022). This may allow the joint activity patterns of On and Off pathways to cover a more versatile stimulus space at the onset of a new fixation than pathways with similar sensitivity profiles, but opposing contrast sensitivity.

## Acknowledgements

This work was supported by the European Research Council (ERC) under the European Union’s Horizon 2020 research and innovation programme (grant agreement number 724822), by the Deutsche Forschungsgemeinschaft (DFG, German Research Foundation) – project numbers 154113120 (SFB 889, project C01) and 432680300 (SFB 1456, project B05), and by the Boehringer Ingelheim Fonds.

